# The architecture of the endoplasmic reticulum is regulated by the reversible lipid modification of the shaping protein CLIMP-63

**DOI:** 10.1101/431106

**Authors:** Patrick A. Sandoz, Robin A. Denhardt-Eriksson, Laurence Abrami, Luciano Abriata, Gard Spreemann, Catherine Maclachlan, Sylvia Ho, Béatrice Kunz, Kathryn Hess, Graham Knott, Vassily Hatzimanikatis, F. Gisou van der Goot

**Affiliations:** Global Health Institute, Life Sciences Faculty, Ecole Polytechnique Fédérale de Lausanne (EPFL), Station 15, CH-1015 Lausanne, Switzerland; Laboratory of Computational Systems Biotechnology, Life Sciences Faculty, Ecole Polytechnique Fédérale de Lausanne (EPFL), Station 15, CH-1015 Lausanne, Switzerland; Institute of Bioengineering, Life Sciences Faculty, Ecole Polytechnique Fédérale de Lausanne (EPFL), Station 15, CH-1015 Lausanne, Switzerland; Brain Mind Institute, Life Sciences Faculty, Ecole Polytechnique Fédérale de Lausanne (EPFL), Station 15, CH-1015 Lausanne, Switzerland; BioEM Facility, Life Sciences Faculty, Ecole Polytechnique Fédérale de Lausanne (EPFL), Station 15, CH-1015 Lausanne, Switzerland

**Keywords:** Endoplasmic reticulum, cellular compartment, S-palmitoylation, ZDHHC6, CLIMP-63/CKAP4

## Abstract

The endoplasmic reticulum (ER) has a complex morphology generated and maintained by membrane-shaping proteins and membrane energy minimization, though not much is known about how it is regulated. The architecture of this intracellular organelle is balanced between large, thin sheets that are densely packed in the perinuclear region and a connected network of branched, elongated tubules that extend throughout the cytoplasm. Sheet formation is known to involve the cytoskeleton-linking membrane protein 63 (CLIMP-63), though its regulation and the depth of its involvement remain unknown. Here we show that the post-translational modification of CLIMP-63 by the palmitoyltransferase ZDHHC6 controls the relative distribution of CLIMP-63 between the ER and the plasma membrane. By combining data-driven mathematical modeling, predictions, and experimental validation, we found that the attachment of a medium chain fatty acid, so-called S-palmitoylation, to the unique CLIMP-63 cytoplasmic cysteine residue drastically reduces its turnover rate, and thereby controls its abundance. Light microscopy and focused ion beam electron microcopy further revealed that enhanced CLIMP-63 palmitoylation leads to strong ER-sheet proliferation. Altogether, we show that ZDHHC6-mediated S-palmitoylation regulates the cellular localization of CLIMP-63, the morphology of the ER, and the interconversion of ER structural elements in mammalian cells through its action on the CLIMP-63 protein.

**Significance Statement:** Eukaryotic cells subcompartmentalize their various functions into organelles, the shape of each being specific and necessary for its proper role. However, how these shapes are generated and controlled is poorly understood. The endoplasmic reticulum is the largest membrane-bound intracellular compartment, accounting for more than 50% of all cellular membranes. We found that the shape and quantity of its sheet-like structures are controlled by a specific protein, cytoskeleton-linking membrane protein 63, through the acquisition of a lipid chain attached by an enzyme called ZDHHC6. Thus, by modifying the ZDHHC6 amounts, a cell can control the shape of its ER. The modeling and prediction technique used herein also provides a method for studying the interconnected function of other post-translational modifications in organelles.

## Introduction

The endoplasmic reticulum (ER) is a complex multifunctional organelle that extends from the nuclear envelope to the cell periphery (1, 2). Based on morphological features, it is classically separated into three distinct subcompartments: the nuclear envelope, the rough ER, and the smooth ER. The rough ER is composed of packed membrane sheets studded with ribosomes and is concentrated in the perinuclear region, while the smooth ER is formed by narrow tubular membranes arranged as a tentacular meshwork that occupies the entire cytoplasm, reaching the plasma membrane. While the major function of the rough ER is considered to be synthesis of proteins targeted to the secretory pathway and the endomembrane system, the smooth ER is thought to be involved in lipid biogenesis, calcium ion storage, and cell detoxification (3).

In the last decade, the mechanisms generating this complex ER architecture have been extensively studied, and several protein families have been observed to control the shape of ER membranes. For instance, ER-sheet formation depends on the protein TMEM170A (4) and on the cytoskeleton-linking membrane protein 63 (CLIMP-63, also called CKAP4) (5-7); ER-sheet flatness depends on kinectin-1 (KTN1) and ribosome binding protein 1 (RRBP1) (2); the width of the ER lumen also involves CLIMP-63 (5-7); the highly curved membranes in the ER tubules and at ER-sheet edges are sustained by ER-curving protein families, such as reticulons (RTNs) and receptor-enhanced expression proteins (REEPs) (8); and, ER tubule fusion is regulated by specific GTPases, the atlastins (ATL) (9). More recently, mathematical models have suggested that ER sheets intrinsically tend to minimize their surface tension energy and form a helicoidal compact arrangement resembling a parking garage to maximize the surface area to volume offered to ribosomes (10). After years of disjointed studies, a somewhat unifying model was finally proposed based on reported observations on the membrane physics and ER-shaping proteins. This model predicts that the specific local ER morphology is regulated by varying the local relative concentrations of various membrane-shaping proteins, reminiscent of a multi-component phase diagram (11).

Even though the mechanisms fundamental to ER shape generation have been heavily studied, it remains unclear how cells control these proteins and, in particular, what governs the interconversion between sheets and tubular structures. In this work, we have studied the effect of a post-translational protein modification, S-acylation, on ER morphology. S-acylation is the addition of a medium-length fatty acid, generally palmitate in a process known as S-palmitoylation, to specific cytosolic cysteine residues of a protein. It is catalyzed by palmitoylacyltransferases (12, 13) and is the only lipid addition that can be reversed, which occurs through the action of acyl protein thioesterases. S-acylation can control the association of soluble proteins with cellular membranes but may also occur on transmembrane proteins to target them to specific membrane subdomains, to induce conformational changes or to drive membrane protein complex formation (14).

Several roles have been assigned to the transmembrane protein, CLIMP-63, which is the only ER-shaping protein for which palmitoylation has been reported (15, 16). It was first described as linking the ER to microtubules through its N-terminal cytosolic tail (17). These studies also indicated that inside the ER, i.e. in the lumen, CLIMP-63 could assemble into dimers and higher order structures, which strongly affect the mobility of the protein in the ER membrane. CLIMP-63 was later proposed to impose the precise width or thickness of the ER lumen by forming dimers in *trans*, i.e. between two CLIMP-63 molecules present in opposing membrane patches “across” the ER lumen (6). CLIMP-63 has also been reported to be present at the plasma membrane, acting as a receptor for various ligands in a tissue-dependent manner (18-21). How these different roles relate to each other, however, and how they are controlled is unknown. CLIMP-63 was reported to undergo two post-translational modifications: phosphorylation, which affects its microtubule binding ability (22), and palmitoylation, which may be mediated by ZDHHC2 (23), a palmitoyltransferase at the plasma membrane (24-26) that may thus play a role in CLIMP-63-mediated signaling at the cell surface.

Here, we show that CLIMP-63 is a preferential target of the ER-localized palmitoylacyltransferase ZDHHC6, and that ZDHHC6 regulates the plasma membrane abundance of CLIMP-63 by controlling its ER exit. We found that ZDHHC6 regulates CLIMP-63 turnover, by stabilizing higher order assemblies, and thus controls its abundance. Strikingly, overexpression of ZDHHC6 drastically affected ER morphology, leading to massive ER-sheet proliferation in a manner dependent on CLIMP-63 and its palmitoylation. Altogether, our results show that ER morphology can be locally and dynamically controlled by the ZDHHC6-mediated palmitoylation of the ER-shaping protein CLIMP-63.

## Results

### The bulk of cellular CLIMP-63 is in a palmitoylated form

We first confirmed that, as reported, CLIMP-63 undergoes palmitoylation, using a resin-assisted capture method, called Acyl-RAC. In this assay, palmitate is removed by hydroxylamine and the freed cysteines are then captured using thiol-reactive beads. The final read out used here was western blot analysis. CLIMP-63 was readily captured using this assay (SI Appendix, Fig. S1). We also tested a panel of reported ER-shaping proteins – ATL2, ATL3, spastin, KTN1, RRBP1, lunapark, and TMEM170A (SI Appendix, Fig. S1a-c, with the validation of the antibodies) – but none were robustly captured. Incubating cells with ^3^H-palmitate led to its incorporation into CLIMP-63, and this incorporation was abolished when Cys-100 was mutated to alanine (SI Appendix, Fig. S1d) (27). For exogenous expression, HA or monomeric red fluorescent protein (RFP) tags were added to the N-terminus of the protein since we found that this recapitulated the behavior of the endogenous protein CLIMP-63, whereas C-terminal tags prevented the ER sheet proliferation induced by CLIMP-63 overexpression (SI Appendix, Fig. S1e).

Since palmitoylation occurs on the cytosolic domain of CLIMP-63 and since this domain is involved in linking CLIMP-63 to the microtubule network, we tested whether palmitoylation and phosphorylation influence one another. The palmitoylation of CLIMP-63 was insensitive to nocodazole-induced microtubule depolymerization or to Taxol-induced microtubule stabilization (SI Appendix, Fig. S2a). It was also insensitive to mutations of the serine phosphorylation sites involved in microtubule binding (SI Appendix, Fig. S2b) (22). Conversely, mutating the CLIMP-63 palmitoylation site did not affect microtubule binding (SI Appendix, Fig. S2c).

We next determined the percentage of cellular CLIMP-63 that is palmitoylated at steady state. We first used an assay where palmitate is exchanged with a PEG molecule, leading to a shift in mass that can be monitored by gel electrophoresis (28, 29). The result showed that the majority of CLIMP-63 is palmitoylated at steady state in our cells (Figure 1a). To gain precision, we developed a variation of the Acyl-RAC method for quantifying free, i.e. non-acylated, cysteines that involves an alkylation step using fluorescent iodoacetamide for a more quantitative detection (Fig. 1b and SI Appendix, Fig. S2d). In addition to Cys-100, CLIMP-63 has only one other cysteine, Cys-126, on the ER lumenal boundary of the transmembrane region. Cys-126 was mutated to alanine to facilitate the quantification of palmitoylation on Cys-100. To analyze CLIMP-63 mutants, we generated a cell line stably expressing an shRNA construct to silence endogenous CLIMP-63 (SI Appendix, Fig S3). Since CLIMP-63 levels influence ER morphology, we titrated the amount of DNA required to reach an endogenous level of CLIMP-63 upon exogenous expression (SI Appendix, Fig. S3c), which was used in all further experiments. Using alkylation with fluorescent iodoacetamide, we could determine that only 12.7 ± 0.05% of the CLIMP-63 population had a free Cys-100 (Figure 1b). Thus, 87% of CLIMP-63 is palmitoylated in our cells. The method can be further modified to probe for acylated, as opposed to free, cysteines leading to the same percentage of palmitoylated CLIMP-63 (SI Appendix, Fig. S2ef).

**Figure 1.**
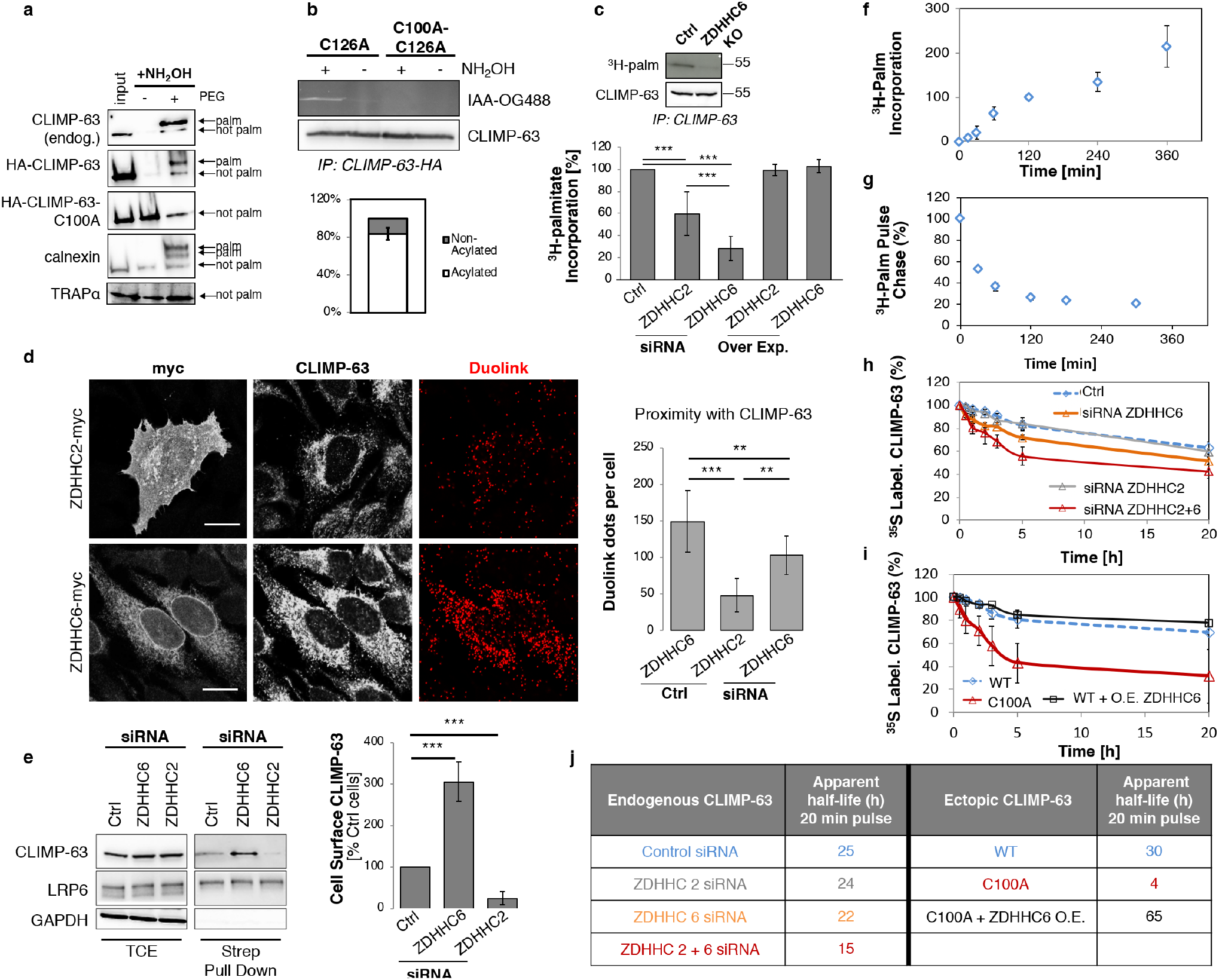
ZDDHC6-mediated CLIMP-63 palmitoylation affects its localization and turnover. **a**. Majority of CLIMP-63 is palmitoylated. Stoichiometry of CLIMP-63 palmitoylation in HeLa cells was followed with the APEGS assay. PEG-5k was used to label endogenous protein or transfected HACLIMP-63, HA-CLIMP-63-C100A, or endogenous Calnexin or endogenous TRAP-alpha (+PEG), in the -PEG the PEG was not added. **b**. Determination of the non-acylated fraction of CLIMP-63. HACLIMP63 C126A or C100A + C126A were treated or not with hydroxylamine (NH_2_OH) and labeled with iodoacteamide-oregon-green-488 (IAA-OG488) as described in SI Appendix, Figure S2d. The amount of acylated CLIMP-63 was determined by comparing plus and minus NH_2_OH (n = 4). Error bar represents standard deviation (SD). **c**. Metabolic labeling with ^3^H-palmitate was performed on ZDHHC6 KO or control HeLa cells for 2 h at 37°C. Bottom panel: ^3^H-palmitate labeling of HeLa cells with control, ZDHHC2, or ZDHHC6 siRNA, or overexpressed ZDHHC2-myc or ZDHHC6-myc for 2 h. Quantification of CLIMP-63 ^3^H-palmitate metabolic labeling with SD (n = 4). ***p < 0.01. **d**. Proximity ligation assay of ZDHHC2 or ZDHHC6 with CLIMP-63 in HeLa control or ZDHHC6 KO cells after cold methanol fixation. Quantification of the proximity ligation assay with SD. (n = 15 each, *p < 0.05; **p < 0.01, ***p < 0.001). **e**. HeLa cell-surface proteins were labeled at 4°C for 30 min with NHS-biotin. LRP6 and GAPDH are blotted as positive and negative controls, respectively. Total cell extract (TCE) = 5% of pulldown volume. Exposure of CLIMP-63 on the right panel is 1.5 orders of magnitude longer than on the left panel. Quantification of CLIMP-63 population at the cell surface with SD (n = 3, ***p < 0.001). **f**. Metabolic labeling with ^3^H-palmitate at 37°C was performed on HeLa cells for different pulse lengths. Quantification of CLIMP-63’s ^3^H-palmitate incorporation showing the average with SD. The signal was normalized so the population at t = 2 h represented 100% (n = 3). **g**. Pulse Chase. After a 2 h ^3^H-palmitate labeling at 37°C, HeLa cells were chased for various times. Quantification of CLIMP-63’s ^3^H-palmitate loss showing the average with SD. The signal was normalized so the initial population at t = 0 represents 100% (n = 5). **h**. Pulse chase. ^35^S Metabolic labeling with a pulse of 20 min was performed on HeLa cells transfected with RNAi CTRL (n = 5) or against ZDHHC2, ZDHHC6, or ZDHHC2 + 6 (all n = 3). The radioactive signal for the population at t = 0 was normalized to 100%. The error bars represent SD**. i**. Pulse chase. Same as (h), but HeLa cells were transfected with either HA-CLIMP-63 WT or C100A, or HA-CLIMP-63 + ZDHHC6-myc together (all n = 3). **j**. Apparent half-lives as extracted from data in (h) and (i).

### CLIMP-63 is a substrate for ZDHHC6, which regulates its plasma membrane targeting

The above experiments indicated that almost 90% of CLIMP-63 is palmitoylated under our experimental conditions. Morphological analyses have reported that CLIMP-63 was predominantly found in the ER (6, 30), as was confirmed later in this study. Combined, these observations imply that ER-localized CLIMP-63 is largely palmitoylated, which in turn suggests that ZDHHC2 (31), given its preferential plasma membrane localization (24,25), cannot be the sole palmitoyl-acyltransferase to modify CLIMP-63. Among the 23 known ZDHHC enzymes, many localize to the ER (32). ZDHHC6 stands out because it has been found to modify various key ER proteins such as calnexin (33), the E3 ligase gp78 (34), and the inositol triphosphate receptor IP3R (35). We therefore investigated its ability to modify CLIMP-63. Incorporation of ^3^H-palmitate into CLIMP-63 was strongly decreased upon CRISPR-Cas9 knockout (KO) of ZDHHC6 (Figure 1c, validation of the KO cells in SI Appendix, Fig. S4a,b) indicating that CLIMP-63 is indeed a target of this enzyme.

We next compared the relative involvements of ZDHHC2 and ZDHHC6. A more pronounced decrease in ^3^H-palmitate incorporation, ~70%, was seen when ZDHHC6 expression was silenced with small interfering RNA (siRNA) than for ZDHHC2 silencing, ~40% (Figure 1c and SI Appendix, Fig. S4c). Overexpression of these enzymes did not affect ^3^H-palmitate incorporation. The lack of increase upon overexpression could be because the majority of CLIMP-63 is already modified at the resting state and thus was not available for modification by ^3^H-palmitate.

Co-immunoprecipitation experiments further indicated that CLIMP-63 could interact with both ZDHHC2 and ZDHHC6 (SI Appendix, Fig. S4d). For a more quantitative evaluation, we performed a dual proximity ligation assay (36) in which the close proximity of two proteins is detected by antibodies coupled to DNA oligonucleotides that are ligated and labeled with fluorescent probes when they are near each other. Interaction could be observed with both enzymes but was considerably more pronounced with ZDHHC6 than with ZDHHC2 (Figure 1d), consistent with the relative abundance of CLIMP-63 in the ER and at the plasma membrane.

As mentioned, a population of CLIMP-63 was reported to act as a cell-surface signaling receptor, indicative of its presence at the plasma membrane (18-21, 37). We investigated whether ZDHHC6 affects the abundance of CLIMP-63 at the plasma membrane using a surface biotinylation assay, wherein all surface proteins were chemically modified, isolated with streptavidin beads, and CLIMP-63 was subsequently identified by western blot. In control HeLa cells, CLIMP-63 was detected at the cell surface, though in minute amounts, probably sufficient for signaling purposes. This population increased three-fold upon ZDHHC6 silencing (Figure 1e). Consistent with this increased surface localization, proximity ligation between CLIMP-63 and ZDHHC2 was higher in ZDHHC6 KO than in control cells (Figure 1d right panel and SI Appendix, Fig. S4e). These observations indicated that ZDHHC6 controls the plasma membrane targeting of CLIMP-63.

Surface biotinylation was also performed on palmitoylation-deficient CLIMP-63 C100A. Intriguingly, it was less abundant than wildtype (WT) CLIMP-63 at the plasma membrane (SI Appendix, Fig. S4f). This observation raises the possibility that palmitoylation by ZDHHC2 could increase the surface residence time of CLIMP-63, i.e. slow down endocytosis and lysosomal targeting, as previously observed for the anthrax-toxin receptor TEM8 (38). To test this hypothesis, we monitored the surface levels of CLIMP-63 upon ZDHHC2 silencing and found that it decreased four-fold (Figure 1e).

Altogether, these findings indicated that the bulk of CLIMP-63 resides in the ER but that a minor population, the size of which is negatively-controlled by ZDHHC6, exits this compartment in a non-palmitoylated state and reaches the plasma membrane where it is retained through palmitoylation by ZDHHC2.

### ZDHHC6 influences CLIMP-63 turnover

We then investigated the dynamics of CLIMP-63 palmitoylation. Measurements of ^3^H-palmitate incorporation as a function of time showed a gradual increase over at least six hours (Figure 1f and SI Appendix, Fig. S4g). We next monitored palmitate turnover by incubating cells with ^3^H-palmitate for 2 hr (the pulse) followed by different incubation times in a label-free medium (the chase). CLIMP-63 was then immunoprecipitated and analyzed by electrophoresis and radiography. ^3^H-palmitate was rapidly lost from labeled CLIMP-63, with almost half of it released within 30 min (Figure 1g and SI Appendix, Fig. S4h). A plateau was reached after two hours, when about 20% of the labeled CLIMP-63 appeared to retain its bound palmitate (Figure 1g). Almost identical kinetics were observed upon ZDHHC2 silencing (SI Appendix, Fig. S4i). Given the low ^3^H-palmitate incorporation signal when silencing ZDHHC6, we could not measure palmitate turnover under these conditions.

We next analyzed the effect of palmitoylation on CLIMP-63 turnover since we previously observed that palmitoylation can drastically affect the half-life of a protein, either increasing it, as for calnexin (39), or decreasing it, as for ZDHHC6 (40). ^35^S Cys/Met metabolic labeling, or pulse-chase, experiments indicated that CLIMP-63 has an apparent half-life (τ_1/2_) of about 25 hours following a 20-min labelling pulse (Figure 1h). Silencing ZDHHC2 did not impact the decay kinetics of CLIMP-63 (Figure 1h) but silencing ZDHHC6 slightly accelerated the apparent decay (τ_1/2_ = 22 hours, Figure 1h and j). Silencing both ZDHHC6 and ZDHHC2 further accelerated the decay (Figure 1h), thus revealing an effect of ZDHHC2 and confirming that ZDHHC6 acts upstream of ZDHHC2.

To further investigate the role of palmitoylation on CLIMP-63 turnover, we monitored the decay of exogenously expressed WT and C100A mutant proteins as well as the effect of overexpressing ZDHHC6. Exogenously expressed CLIMP-63 showed similar decay kinetics to the endogenous protein (Figure 1i). The overexpression of ZDHHC6 led to an increase in the apparent stability of CLIMP-63 (Figure 1i). Conversely, mutating the palmitoylation site (C100A) led to accelerated degradation, with an apparent half-life of only four hours (Figure 1i-j). The difference in CLIMP-63 stability in ZDHHC6-ZDHHC2 double-silenced cells and CLIMP-63-C100A might be due to the residual presence of the palmitoyl-acyltransferases in the siRNA-treated cells.

Taken together, these results indicate that the palmitoylation of CLIMP-63 is dynamic and that palmitoylation strongly affects the turnover rate of the protein.

### Understanding CLIMP-63 assembly and trafficking through mathematical modeling

The above analyses, combined with knowledge from literature, indicate that the majority of CLIMP-63 resides in the ER with small amounts at the plasma membrane, that in each of these compartments it could be unmodified or contain the palmitoyl modification, and that ZDHHC6 levels modulate the plasma membrane abundance of CLIMP-63. Our data also indicated that while the apparent turnover of palmitate was rapid (Fig. 1g), the vast majority of cellular CLIMP-63 at any given time was in a palmitoylated state (Fig. 1a-b). To understand how these findings fit together, we generated a computational representation of the system and made use of mathematical modeling.

The model was initially composed of five CLIMP-63 species: non-palmitoylated monomer in the ER (M^0^_ER_, the 0 superscript indicates that the palmitoylation site is free), palmitoylated monomer in the ER (M^1^_ER_, the 1 superscript indicates that the palmitoylation site is occupied), the same molecules but at the plasma membrane (PM), M^0^_PM_ and M^1^_PM_, and a nonpalmitoylated intermediate species representing transport from the ER to the PM. Although pulse-chase experiments were accurately reproduced, we could not find reasonable parameter values that would sensibly fit the subcellular distribution of CLIMP-63 (SI Appendix, Fig. S5ab), instead simulating CLIMP-63 as equally abundant at the plasma membrane and in the ER. This suggested that the mechanistic network model would require modification.

One of the first plausible modifications to the structure of the model would take into account the reported observations that CLIMP-63 can form dimers and possibly higher order oligomeric structures (5, 41, 42). Since the information currently available on oligomerization of CLIMP-63 is insufficient to differentiate dimerization in *cis* and in *trans*, the two where lumped in our conceptual model. Palmitoylation states of these dimers was however taken into account, leading to three species, D^0^_ER_, D^1^_ER_ and D^2^_ER_, corresponding to non-palmitoylated, single palmitoylated, and double palmitoylated dimers, respectively (Figure 2e). The same dimerization constant was imposed for M^0^_ER_ and M^1^_ER_. Integrations of higher order multimers of CLIMP-63 were also tested but did not change the behavior of the system, so the dimer structure was retained according to the principle of parsimony (SI Appendix, Fig. S5). Since our experimental data did not reveal any cross-talk between palmitoylation and phosphorylation/microtubule binding, the latter was not included in the mechanistic framework, which was designed to estimate parameters such as protein turnover, palmitoylation and depalmitoylation rates.

**Figure 2.**
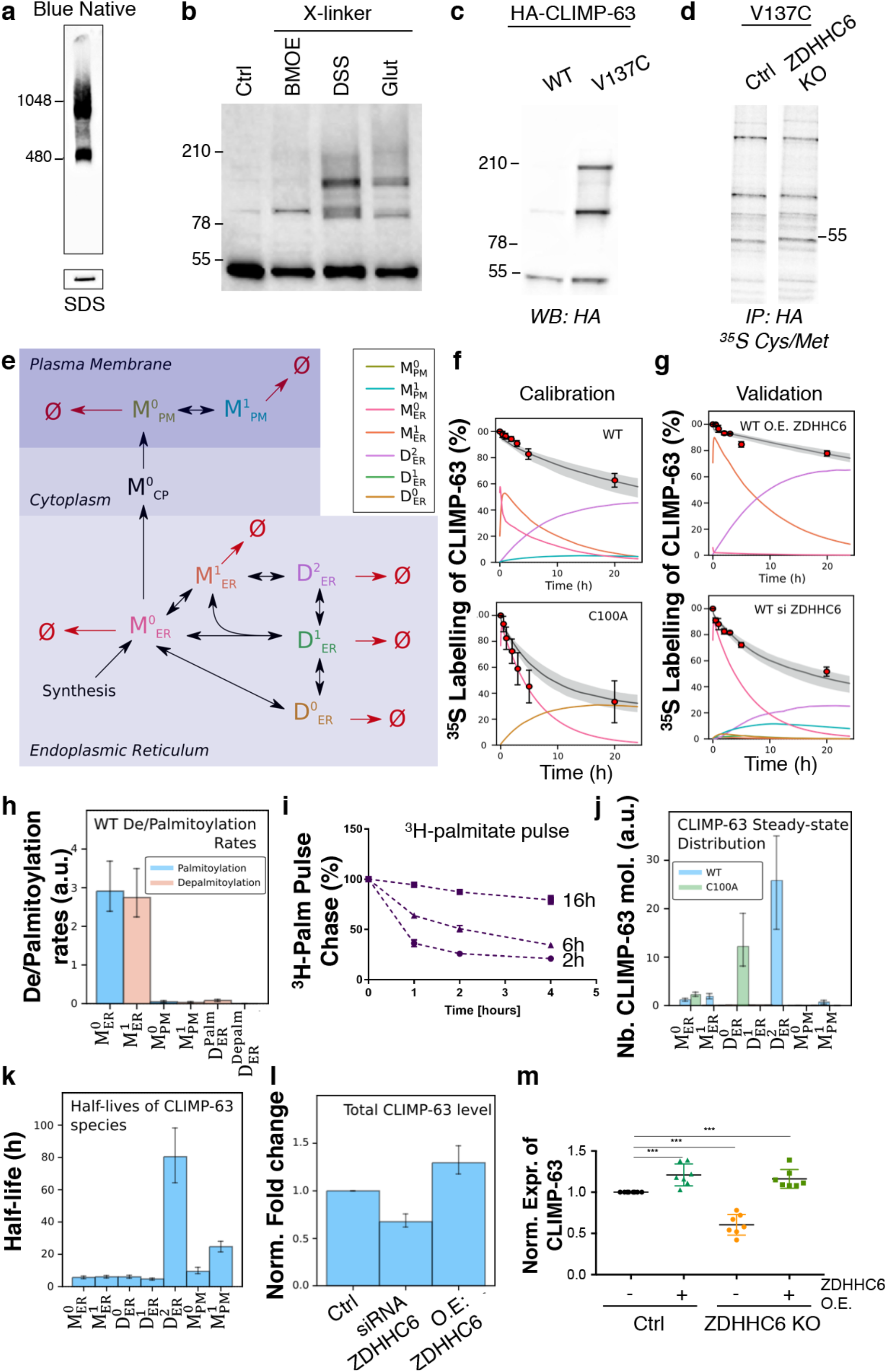
Modeling of CLIMP-6 palmitoylation dynamics. **a**. A post-nuclear fraction of HeLa cells was prepared and ultracentrifuged for 1 h at 45,000 rpm. The membrane fraction was retrieved and separated into two samples migrated either on Blue Native gel or SDS-PAGE. **b**. Endogenous CLIMP-63 in HeLa cells was crosslinked 5 min at room temperature with either 1mM BMOE, 1mM DSS or 0.025% Glutaraldehyde. Total cell extracts were suspended in non-reducing SDS-PAGE sample buffer, run on a 4–20% SDS-PAGE Tris-Glycine gel and revealed with anti-rabbit CLIMP-63 antibodies. **c**. HeLa cells were transfected with HA-CLIMP-63 WT or HA-CLIMP-63 V137C for 24 hours. Total cells extracts were migrated on non-reducing conditions on a 4–20% SDS-PAGE Tris-Glycine gel and revealed with anti-HA antibodies. **d**. Autoradiography of HeLa cells transfected with HA-CLIMP-63 WT or HA-CLIMP-63 V137C for 24 hours, ^35^S Cys-Met radiolabeled for 20 min pulse, immunoprecipitated using HA-beads and loaded on SDS-PAGE under non-reducing conditions. **e**. Schematic description of CLIMP-63 species and localizations. **f**. Calibration and **g**. validation of the model output (from Fig. S2). The solid grey line is the median of 100 model simulations; the shaded grey interval is defined by the 1st and the 3rd quartile of the 100 model simulations (see SI Appendix,). Each species is color coded. **h**. Predicted palmitoylation and depalmitoylation rates for CLIMP-63 species. Rates are lumped together for dimer species. ie. D^palm^_ER_ corresponds to the sum of D^0^_ER_ and D^1^_ER_ palmitoylation rates. **i**. Pulse chase. After either 2 h, 6 h, or 16 h of metabolic labeling with ^3^H-palmitate at 37°C, the HeLa cells were washed with complete medium and lysed at the various time points. The graph represents the ^3^H signal quantification of endogenous CLIMP-63 with error bars as SD (n = 3). **j**. Steady-state distribution of CLIMP-63 species as predicted for CLIMP-63 WT and C100A mutant. Error bars represent the first and third quartile through the simulation of 100 models. **k**. Simulated *bona fide* half-lives of CLIMP-63 species. **l**. *In silico* prediction of the total level of CLIMP-63 under control conditions, ZDHHC6 silencing, or ZDHHC6 overexpression. **m**. Expression of endogenous CLIMP-63 in HeLa cells control, in Hela cells KO for ZDHHC6 and in HeLa ZDHHC6 KO cells recomplemented with ZDHHC6. Endogenous CLIMP-63 of 10µg of total cell extracts were revealed by Western Blotting with rabbit anti-Climp-63 and anti-rabbit alexa-488 antibodies, detected with FUSION Camera and quantified with BIO1D software (n = 8, error bars rereesent SD).

A subset of the experimental data (from Figure 1 and SI Appendix, Fig. S5e) was used to calibrate the model (Figure 2f and SI Appendix, Fig. S5f). A heuristic optimization method generated a population of models consistent with the calibration experiments. We chose the 100 sets of parameters that best fitted the experimental data. These sets of parameters were subsequently used to predict the results of a second set of experiments. As shown in Figure 2g and SI Appendix, Figure S5g, the results fit the predictions. Importantly, with the introduction of dimerization in the ER, the model predicted that the majority of CLIMP-63 is retained in the ER (see below) and suggested that dimerization is the major ER-retention mechanism of CLIMP-63. Since the mathematical equations both fitted a subset of data and predicted another set of data, it was considered as accurately capturing the overall system and appropriate to interpret our data without the necessity to increase the complexity.

Considering the predicted importance of dimerization, we first verified that CLIMP-63 can indeed form dimers. We analyzed CLIMP-63 by Blue-Native PAGE, a method for monitoring protein complexes by electrophoresis. CLIMP-63 migrated with an apparent molecular weight of 480 kDa (Fig. 2a). While this observation does indicate that CLIMP-63 is mostly non-monomeric, it is difficult to draw conclusions regarding multimerization states since conformation and shape also strongly affect protein migration in such gels. We therefore performed experiments with three types of cross linkers: BMOE, a short (8Å) sulfhydryl-to-sulfhydryl crosslinker; DSS, a sulfo-NHS ester that reacts rapidly with any primary amine; and glutaraldehyde, for indiscriminate cross-linking. All three reagents revealed the presence of dimers, while DSS and glutaraldehyde indicated that CLIMP-63 could also associate into higher order structures (Figure 2b), which could include hetero-oligomers. To further confirm the existence of CLIMP-63 dimers, we performed co-immunoprecipitation experiments with HA- and RFP-tagged CLIMP-63 combined with ^35^S Cys/Met metabolic labeling and found that dimers did form, shortly after synthesis, even for the C100A mutant (SI Appendix, Fig. S5c). To get a more accurate estimation of the abundance of dimers and higher order assemblies, we constructed a mutant of CLIMP-63 with a cysteine residue in the ER luminal domain to generate disulfide-bonded oligomers. The entire 472-residue CLIMP-63 domain in the ER lumen is predicted to form a coiled-coil, with two or three breaks (Multicoil2: http://cb.csail.mit.edu/cb/multicoil2/cgi-bin/multicoil2.cgi). We chose to mutate a hydrophobic residue close to the transmembrane domain likely to be involved in monomer-monomer contact, Val-137, to cysteine in order to trap dimers through disulfide bond formation. The electrophoresis migration pattern of the V137C mutant under non-reducing conditions indeed revealed the formation of dimers as well as higher molecular weight forms (Fig. 2c) that must involve other cysteines, either within CLIMP-63 or in an interacting protein. Dimerization of V137C CLIMP-63 was observed rapidly after synthesis as revealed by ^35^S Cys/Met metabolic labeling, both in control cells and in ZDHHC6 KO cells (Fig. 2d), confirming that palmitoylation is not necessary for multimerization. Altogether this analysis indicated that the majority of CLIMP-63 rapidly forms dimers or higher order structures. The observed migration patterns also indicated that CLIMP-63 multimerization is complex. Its understanding will require structural determination of the luminal domain, which probably simultaneously forms both parallel and antiparallel coiled-coil domains at the N-terminal and C-terminal end, respectively, with CLIMP-63 molecules in the same ER membrane plane and with CLIMP-63 molecules in the opposing ER membrane “across” the lumen.

The model allowed the deconvolution of experimental curves into the evolution of the individual species as a function of time. This indicated that when performing a 20-min labeling pulse with ^35^S-Cys/Met, newly synthesized M^0^_ER_ was rapidly converted to M^1^_ER_, which itself was rapidly decayed to partially give rise to D^2^_ER_, the only significant ER species remaining at the 20-hour chase time (Figure 2f, top panel). While the model predicted that monomers underwent rapid palmitoylation in the ER, it also indicated that monomers underwent significant depalmitoylation (Figure 2h). This balance was predicted to result in a 1.6x excess of M^1^_ER_ over M^0^_ER_, increasing the probability of dimerization between M^1^_ER_ but not excluding that M^0^_ER_ can dimerize, as confirmed by dimerization of the V137C mutant in ZDHHC6 KO cells (Fig. 2d) and of HA-CLIMP-63 C100A with RFP-CLIMP-63 C100A (SI Appendix, Fig. S5d).

Interestingly, the palmitoylation and depalmitoylation fluxes of the dimers were predicted to be very low (Figure 2h), leading us to analyze what was being measured during the pulse-chase experiments shown in Fig. 1g. According to the model, after two hours of ^3^H-palmitate labeling, the labeled population was composed of 77% M^1^_ER_ and only 15% D^2^_ER_. M^1^_ER_ was subsequently depalmitoylated while D^2^_ER_ was not, explaining the plateau that the depalmitoylation curve reached at long chase times (Fig. 1g). As for any labeling experiment, the species distribution at the end of the pulse was influenced by the length of the pulse. The model would therefore predict that increasing the length of the pulse would lead to more dimer formation, which would in turn lead to slower apparent depalmitoylation kinetics during the chase (SI Appendix, Fig. S5g). This could be validated experimentally; indeed, depalmitoylation occurred at far slower apparent rates when pulse times were longer (Fig. 2i). Thus, experiments and modeling indicated that dimerization protects CLIMP-63 from depalmitoylation.

We next used the model to infer the half-lives of the different species in order to better describe how CLIMP-63 stability may be regulated. As opposed to experiments that provide the apparent half-lives of mixed protein populations, models can predict half-lives of individual species. All ER-localized CLIMP-63 species were predicted to have similar half-lives, around five hours, with the notable exception of D^2^_ER_ whose half-life was predicted to be above 80 hours (Figure 2k). To validate this prediction, we sought a method that would allow us to estimate the half-life of the majority of the cellular CLIMP-63 population, as opposed to ^35^S-Cys/Met labeling that monitored a newly synthesized population that gradually converted to different species as a function of time. For this, we performed fluorescent pulse-chase experiments using a fusion protein of CLIMP-63 with a SNAP tag. This tag allows labeling of a fully folded protein and can be used to measure its half-life (39). No SNAP-CLIMP-63 degradation was observed over a 24 h period, confirming that the most abundant species has a very long half-life (SI Appendix, Fig. S5h).

Prediction of the species distribution indicated that the palmitoylated dimer was the most abundant CLIMP-63 species (Figure 2j), consistent with its long half-life and very slow depalmitoylation. The overall abundance of CLIMP-63 was predicted to depend on ZDHHC6 activity in cells, whose overexpression was predicted to increase the overall CLIMP-63 level by 30% (Figure 2k) due to the accelerated palmitoylation of monomers and the subsequent production of palmitoylated dimers (SI Appendix, Fig. S5i, magenta curves). ZDHHC6 silencing was predicted to decrease CLIMP-63 levels by 32% (Figure 2l). These predictions could be experimentally validated since the expression of CLIMP-63 was 30% lower in ZDHHC6 KO cells and about 20% higher when overexpressing ZDHHC6 (Fig. 2m).

Finally, consistent with experimental observations, the model predicted that ZDHHC6 controlled the concentration of the ER-exit-competent species and, thereby, the plasma membrane population. At the surface, ZDHHC2-mediated palmitoylation of CLIMP-63 was predicted to delay degradation, presumably by delaying targeting to lysosomes. Thus, the plasma membrane population of CLIMP-63 that is involved in signaling events appears to be controlled negatively by ZDHHC6 and positively by ZDHHC2.

In summary, the synergistic cycle of experiments, model development, model-based analysis of experimental data and simulations support the conclusions that dimerization and palmitoylation jointly, but not alone, drastically stabilize CLIMP-63, that dimerization prevents depalmitoylation, and thus that the palmitoylated multimers are far more abundant than monomers in the cells. The joint importance of palmitoylation and dimerization was also apparent when performing a global sensitivity analysis (SI Appendix, Fig. S5j). The kinetic properties of both the palmitoylation of CLIMP-63 dimer and monomer (SI Appendix, Fig. S5j) were indeed among the most important parameters for an accurate calibration of the model.

### CLIMP-63 palmitoylation regulates ER morphology

CLIMP-63 overexpression was reported to induce ER-sheet proliferation (5, 6). Since our data-driven mathematical modeling indicated that the abundance of CLIMP-63 was strongly influenced by palmitoylation, we investigated whether this lipid modification played a role in shaping the ER. We choose to first modulate depalmitoylation kinetics. The model predicted that a decrease in palmitate turnover would lead to an accelerated accumulation of dimers and an increased apparent half-life when performing ^35^S Cys/Met pulse-chase experiments (Fig. 3a). In membrane proteins, palmitoylation sites are often found in close proximity to the transmembrane domain and often occur in pairs, such as in the anthrax toxin receptor TEM8 (38) or the ER chaperone calnexin (33), where dual site modification essentially prevented depalmitoylation (39). We thus tested if inserting a second cysteine right next to Cys-100, which is five residues away from the transmembrane domain, would affect the palmitate turnover rate of CLIMP-63 monomers. This insertion is unlikely to have structural consequence since the cytosolic tail of CLIMP-63 is predicted to be disordered (https://iupred2a.elte.hu/). As predicted, the depalmitoylation of CLIMP-63-CC was drastically slower after a two-hour ^3^H-palmitate pulse than observed for WT, with an almost 10-fold increase in the apparent half-life of bound palmitate (Figure 3b). The palmitate decay plateaued at 40% (Figure 3b), consistent with the predicted higher percentage of palmitoylated dimers at the end of the pulse compared to WT CLIMP-63. Finally, consistent with the effect of palmitoylation on CLIMP-63 turnover, the apparent half-life of CLIMP-63-CC was longer than that of WT (Figure 3c). We next analyzed the effect of CLIMP-63-CC expression on ER morphology. We observed a striking densification of perinuclear ER-sheets, as quantified over some 100 CLIMP-63-CC-expressing cells randomly chosen over three independent experiments (Figure 3d). This observation indicated that the dynamics of CLIMP-63 palmitoylation strongly influence ER morphology.

**Figure 3.**
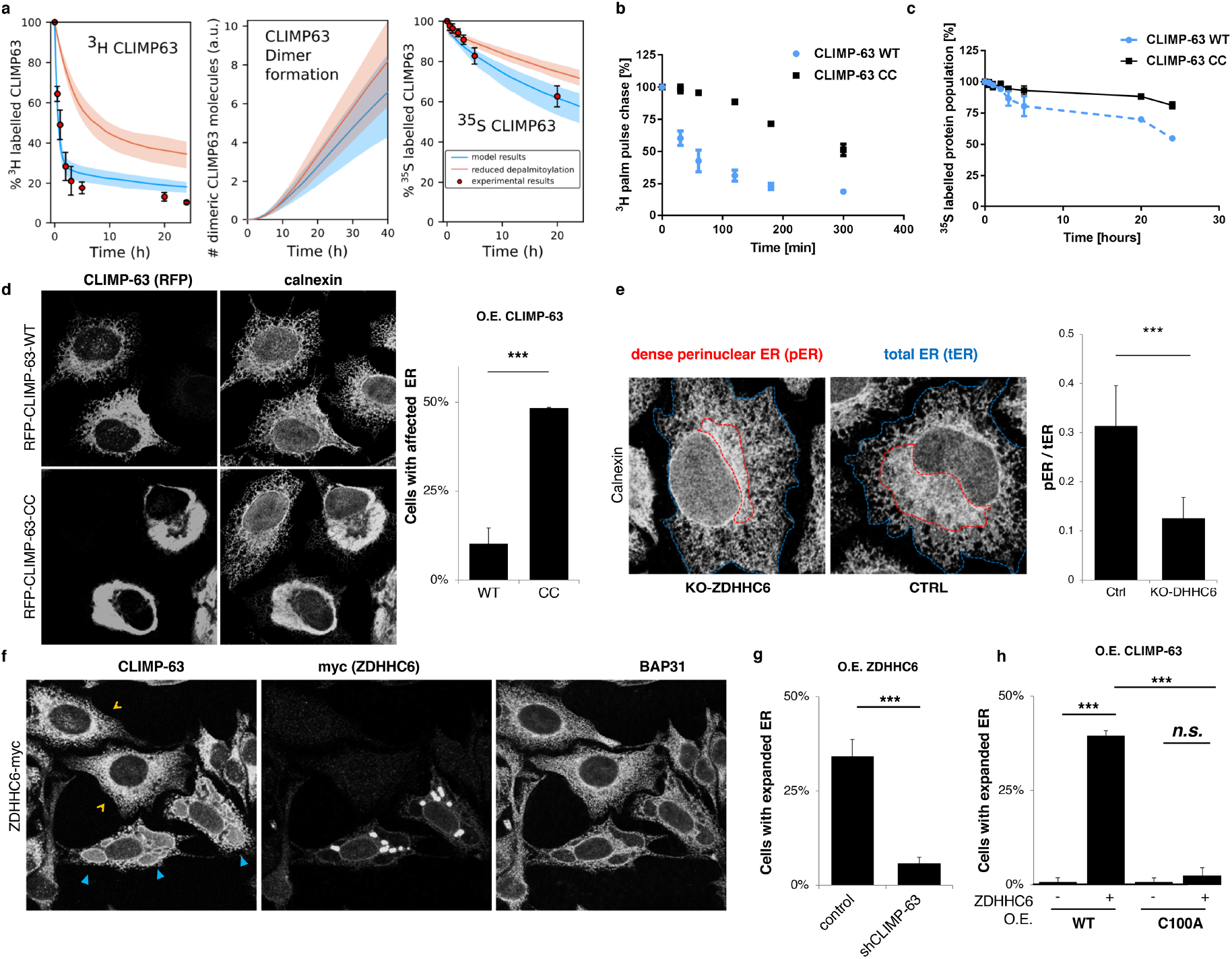
ZDHHC6-mediated CLIMP-63 palmitoylation induces ER-sheet proliferation. **a**. Computational simulations of CLIMP-63 depalmitoylation (left), dimer formation (middle), and protein stability (right) upon slower depalmitoylation kinetics. Median shown by solid lines, 1^st^ and 3^rd^ quartile by shaded interval. **b**. Pulse chase: ^3^H-palmitate pulse (2 h)-chase of HeLa cells transfected with FLAG-CLIMP-63 WT or CC mutant (n = 3). The plot represents the quantification with average and standard deviation (SD). The signal was normalized to that at t = 0. **c**. Pulse chase: ^35^S metabolic pulse (20 min)-chase of HeLa cells transfected with HA-CLIMP-63 WT or CC (n = 3). The plot represents the quantification with average and SD. The signal was normalized so the initial population at t = 0. **d**. HeLa cells stably expressing a shRNA against CLIMP-63 were recomplemented for 48 h with either RFPCLIMP-63 WT or CC and analyzed by immunofluorescence. Calnexin and TRAPα were used as ER markers. Quantification represents the percentage of cells with higher perinuclear density of the ER (n = 3 with each > 30 randomly chosen cells, ***p < 0.01). **e**. HeLa CRISPR-Cas9 ZDHHC6 KO cells or control cells were analyzed by immunofluorescence. The surfaces of the dense perinuclear ER (pER; red) as well as the total ER (tER; blue) were measured using ImageJ. The bar plot represents the ratio of pER/tER for the control and KO-ZDHHC6 cells (32 different randomly chosen cells were measured for each condition; ***p < 0.001). **f**. HeLa cells were transfected for 48 h with ZDHHC6-myc and analyzed by immunofluorescence. Blue arrows highlight the dense perinuclear ER sheets while orange arrows show non-transfected cells. **g**. As in (f) using HeLa cells stably expressing a shRNA against CLIMP-63. Quantification of the proportion of cells overexpressing ZDHHC6-myc with a dense perinuclear region expansion in the absence or not of CLIMP-63 (n = 3). ZDHHC6 in control (CTRL): 129 cells; ZDHHC6 in shCLIMP-63: 226 cells. ***p < 0.01. **h**. HeLa cells transfected for 48 h with ZDHHC6-myc and CLIMP-63 WT or C100A mutant were analyzed by immunofluorescence. Quantification of cells with or without induced ER-sheets (n = 3). Cells with expanded ER were normalized to the total number of cells. HA-CLIMP-63-WT: 137 cells, HA-CLIMP-63-WT + ZDHHC6: 79 cells, HA-CLIMP-63-C100A: 145 cells, HA-CLIMP-63-C100A + ZDHHC6: 182 cells. ***p < 0.01, n.s.: non-significant.

We next modulated ZDHHC6 expression levels. In ZDHHC6 KO HeLa cells, the ER appeared more tubular than in control cells and showed a decreased density in the perinuclear region (Figure 3e and SI Appendix, Fig. S6b). Conversely, overexpression of ZDHHC6 in HeLa and U2OS cells clearly induced severe alterations in ER morphology (SI Appendix, Fig. S6cd). Importantly, ZDHHC6 overexpression did not induce ER stress, as shown by qPCR analysis of the mRNA levels of major ER stress mediators BiP, Ire1, PERK, and ATF6 (SI Appendix, Fig. S6e). This overexpression caused ZDHHC6 to accumulate into dot-like structures when visualized by fluorescence microscopy, which corresponded to organized smooth endoplasmic reticulum (OSERs) as revealed by electron microscopy (Fig. 3f and SI Appendix, Fig. S7) (40, 46). These ER alterations were specific to ZDHHC6 since the overexpression of other ER-residing ZDHHC enzymes, such as of ZDHHC24, did not induce these morphological changes (SI Appendix, Fig. S8). Thus, ZDHHC6 expression levels directly influence ER architecture, as further analyzed below.

We next investigated whether the effect of ZDHHC6 on ER morphology was mediated by CLIMP-63. Remarkably, ZDHHC6 overexpression had no effect on ER morphology in CLIMP-63 knockdown cells (Figure 3g and SI Appendix, Fig. S8a). Expression of exogenous, small hairpin RNA (shRNA)-resistant, WT CLIMP-63 restored the ZDHHC6-induced ER proliferation phenotype, whereas expression of the palmitoylation-deficient C100A mutant did not (Figure 3h and SI Appendix, Fig. S8b). Thus, ER-sheet proliferation induced by ZDHHC6 overexpression depends on palmitoylation of CLIMP-63.

To investigate the impact of CLIMP-63 palmitoylation on ER morphology at a higher resolution, we performed electron microscopy analysis. We expressed RFP-tagged WT in cells stably transduced with CLIMP-63 shRNA. These cells were analyzed by correlation microscopy: first by identifying transfected cells by fluorescence microscopy and subsequently analyzing these cells by transmission electron microscopy. We could not tag ZDHHC6 with GFP because it led to functional alterations. CLIMP-63 itself was therefore tagged at the N-terminus with RFP. As expected, long ER-sheets were observed in CLIMP-63 shRNA cells expressing RFP-CLIMP –63 (SI Appendix, Fig. S10). We next overexpressed ZDHHC6-myc. High ZDHHC6 expressing cells could be identified by the presence of dark dot-like structures, the OSERs, in the RFP-CLIMP-63 staining pattern (SI Appendix, Fig. S10 according to Figure 3h and SI Appendix Fig. S7). We next imaged the cell volume using focused ion beam scanning electron microscopy (FIBSEM). This technique provided serial images with near isotropic voxels from which we reconstructed the ER (Figure 4a-d and SI Appendix, Movie S1 to S3). The ER sheets appeared as a loose, stratified matrix containing multiple openings between layers. Upon ZDHHC6 overexpression, the layering pattern appeared denser with less openings in the ER sheets, and multiple convolutions of the membrane (Figure 4d, Movie S3).

**Figure 4.**
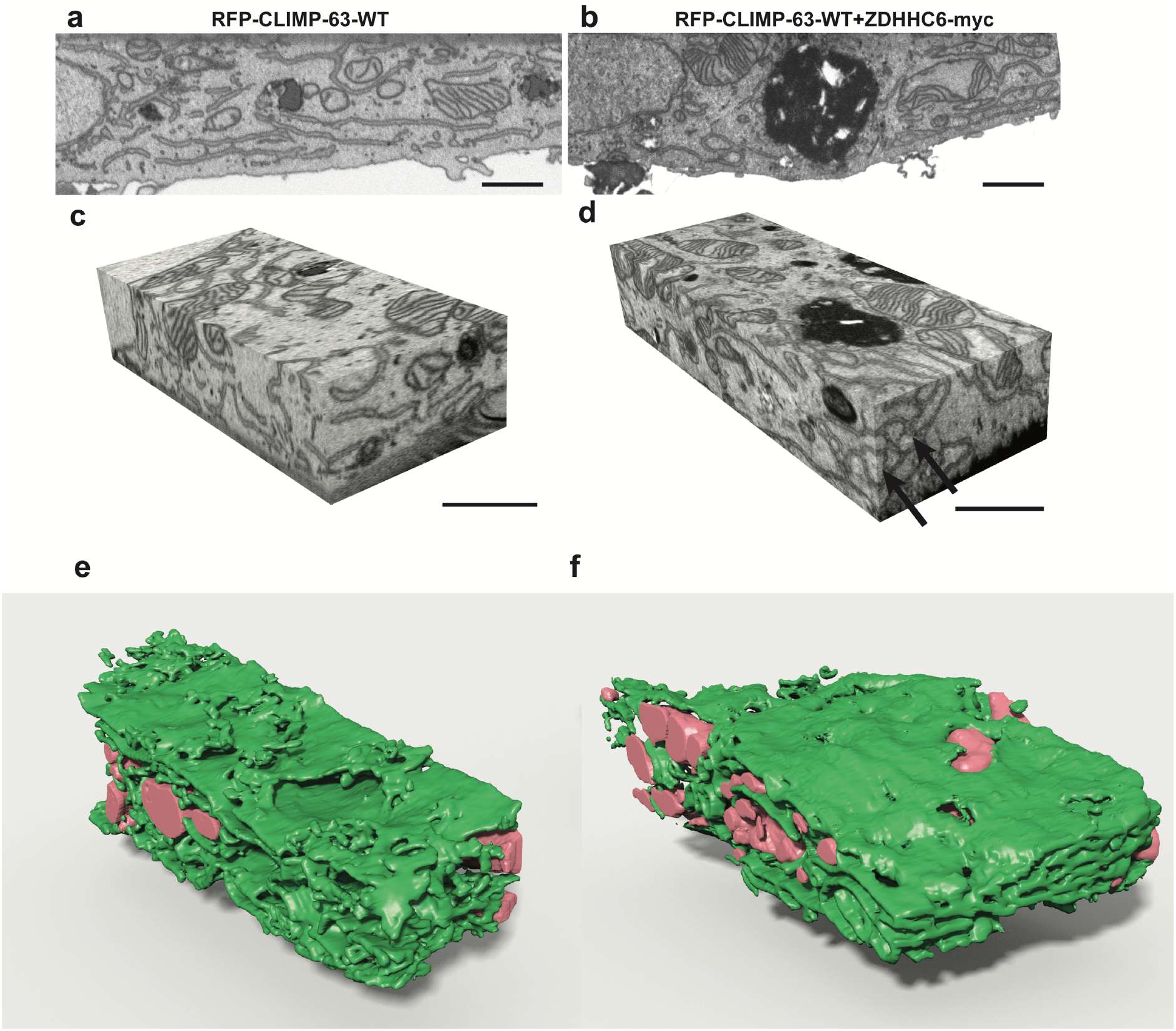
ZDHHC6 induced ER changes. FIBSEM was used to 3D image the ER in RFP-CLIMP-63 under control conditions (**a**) or upon co-overexpression of ZDHHC6 (**b**). **c-d**. FIBSEM image stacks, with near isotropic voxels, show the convoluted branching pattern of the ER. In (**d**), numerous closed loops of membrane can be seen in the two imaging planes (arrows). Reconstructions of ER (green) show its conformation in cells under control conditions (**e**) or upon ZDHHC6-myc overexpression (**f**), with the reconstructed mitochondria (pink) and nuclear membrane (yellow).

The FIBSEM images and the 3D reconstruction revealed an increase in continuity and a densification of the ER sheets upon ZDHHC6 overexpression in WT CLIMP-63-expressing cells (Fig. 4, Movie S3). In order to quantify these visual changes, we employed persistent homology, a construction from the mathematical field of applied algebraic topology (for thorough and mathematically precise introductions, please refer to previous studies [47-51]). Persistent homology can track the appearance and disappearance of features — spherical cavities (in degree-2) and loops (degree-1) — in data across a range of scales (SI Appendix, Fig. S11–13). We summarize persistent homology as a diagram with a point for each feature. The point’s horizontal coordinate encodes the feature’s appearance, and its vertical coordinate encodes its disappearance (in the artificial case illustrated in (SI Appendix, Fig. S11a), the diagram would contain a single point [b, e]). Persistent homology of both degree-1 and degree-2 (SI Appendix, Fig. S11b-f) confirms the change in ER topology upon ZDHHC6 overexpression, with a loss of ER fenestration and an increase in sheet density.

## Discussion and Conclusions

This study investigated the role of the reversible post-translational modification S-palmitoylation on the complex and dynamic architecture of the ER. ER morphology has been previously shown to be dependent on the N-myristoylation of the protein Lunapark (52). Because myristoylation is irreversible, however, it is unlikely to dynamically regulate ER membrane rearrangements. Independent studies showed that CLIMP-63 can be palmitoylated and that it is involved in ER sheet formation. Because the reversible nature of palmitoylation makes it ideal for mutable systems, we analyzed the palmitoylation status and dynamics of CLIMP-63 to determine whether it influences ER-sheet formation.

Based on the experimental determination of palmitoylation, depalmitoylation, and protein turnover combined with data-driven mathematical modeling and prediction, we propose the following outlook on CLIMP-63 involvement in ER-shaping and plasma membrane signaling. As with most transmembrane proteins, CLIMP-63 is synthesized by ribosomes that dock onto the translocon on the ER membrane, generating monomeric CLIMP-63 (M^0^_ER_). CLIMP-63 does not harbor any classical ER retention signals, allowing M^0^_ER_ to exit the ER and reach the plasma membrane. CLIMP-63 can be retained in the ER through two independent mechanisms: S-palmitoylation or dimerization. M^0^_ER_ is highly susceptible to palmitoylation by ZDHHC6, generating M^1^_ER_, which can either be rapidly depalmitoylated by an acyl-protein thioesterase that remains to be identified (depalmitoylation cannot be spontaneous and requires a thioesterase), or it can dimerize. As opposed to monomers, dimers do not undergo significant depalmitoylation and have an ~16-times longer half-life than all other ER-localized CLIMP-63 species. In the absence of palmitoylation, dimerization also occurs, leading to ER retention. Thus, the vast majority of CLIMP-63 remains in the ER and is almost exclusively in the palmitoylated multimeric form.

Palmitoylation controls the residence time of CLIMP-63 in both the ER and at the plasma membrane. Non-palmitoylated CLIMP-63 monomers can be transported out of the ER to the cell surface, with the amount exiting dependent on ZDHHC6. Experiments indeed indicate that the amount of CLIMP-63 at the cell surface, where it is required in lower numbers due to its involvement in signaling (18-21, 37), can vary several fold in response to changes in ZDDHC6 activity, with less palmitoylation by ZDDHC6 allowing for an increased presence of CLIMP-63 at the plasma membrane. It always, however, remains a very minor proportion of the total cellular CLIMP-63 population. At the plasma membrane, CLIMP-63’s abundance is again under the control of palmitoylation, this time through the action of ZDHHC2 (25, 31, 53). M^1^_PM_ has a longer predicted half-life than M^0^_PM_, thus suggesting that ZDHHC2-mediated palmitoylation increases the surface residence time of CLIMP-63, as reported for the anthrax toxin receptor TEM8 (38). Altogether, the present study shows that the distribution of various species of CLIMP-63 as well as their localization in the cell are influenced by ZDHHC6-mediated palmitoylation in the ER and ZDHHC2-mediated palmitoylation at the plasma membrane.

The most striking observation is that CLIMP-63 palmitoylation dramatically affects ER morphology. Palmitoylation was perturbed by overexpression or KO of ZDHHC6, expression of the palmitoylation-deficient CLIMP-63 C100A mutant, and expression of the CLIMP-63-CC mutant that harbors two neighboring palmitoylation sites leading to slower palmitate turnover on monomers. Analyses by fluorescence and electron microscopy showed that proper control of the ER morphology requires well-tuned palmitoylation. Severe alterations were indeed caused by the absence of CLIMP-63 palmitoylation, decreased depalmitoylation kinetics, or excessive palmitoylation. In particular, excessive palmitoylation of CLIMP-63 led to the proliferation of ER sheets, which increased in size and became extremely smooth, without fenestrations, as revealed by FIBSEM microscopy. A sort of phase diagram has been proposed to explain changes in ER structural elements based on variations in other known ER-shaping proteins (11). It is tempting to speculate that local concentrations of CLIMP-63 species, particularly in peripheral ER sheets surrounded by tubules, may reside near phase transitions. Consequently, moderate ZDHHC6-induced variations in CLIMP-63 dimers may trigger pronounced tubule-to-sheet conversions. In addition, palmitoylation may influence local membrane curvature as recently reported for the influenza virus hemagglutinin (54).

Most early studies on CLIMP-63 were focused on its ability to bind microtubules (17) through the phosphorylation of three serines in the cytosolic domain of CLIMP-63 (22), thereby affecting the microtubule network. Here we show that the zDHHC6-mediated palmitoylation of CLIMP-63, while not influenced by microtubule binding, strongly influences ER morphology and in particular the abundance of ER-sheets. The activity of zDHHC6, which controlled by the upstream palmitoyltransferase zDHHC16, varies in a tissue dependent manner (40) and future studies will elucidate how this impact ER structure. Palmitoylation may also occur for other ER-shaping proteins in addition to CLIMP-63, in particular tubule generating protein, which would allow a coordinated generation of tubules and sheets.

## Materials and Methods

### Cell culture, transfections, immunoprecipitation, and western blotting

All HeLa cells were cultured in MEM Eagle (Sigma, US) complemented with 10% FCS (PAN Biotech, D), 1% Pen/Strep, 1% L-Glutamine, and 1% MEM NEAA (all from Gibco, US). HeLa cells are not on the list of commonly misidentified cell lines maintained by the International Cell Line Authentication Committee. Our cells were authenticated by Microsynth (CH), which revealed 100% DNA identity with ATCC®CCL-2™. They were mycoplasma negative as tested on a trimestral basis using the MycoProbe Mycoplasma Detection Kit CUL001B. For transfection, the cells were dissociated using trypsin and plated in tissue culture dishes (Falcon, US). After 24 h, the medium was changed and the cells were transfected using Fugene for plasmids (Promega, US) or Interferrin (Polyplus, F) for silencing with siRNA. The cells were incubated for 24 h to 48 h (for plasmids) or 72 h (for siRNA) before performing experiments.

For immunoprecipitation and western blotting the cells were lysed on ice for 30min with lysis buffer (500 mM Tris–HCl pH 7.4, 2 mM benzamidine, 10 mM NaF, 20 mM EDTA, 0.5% NP40 and a protease inhibitor cocktail (Roche, CH)). The lysate was then centrifuged at 4°C for 3 min at 5000 rpm and the supernatant retrieved. For immunoprecipitation, the lysate was pre-cleared using Sepharose G-beads without antibody for 30 min at 4°C before immunoprecipitating (with G-beads and antibody) turning on a wheel overnight at 4°C. The beads were then washed 3x with lysis buffer before adding sample buffer (4x) including beta-mercaptoethanol. The samples were boiled 5 min at 95°C and vortexed before loading and migration on 4–12% or 4–20% Tris-glycine SDS-PAGE gels. After 2 h at 120 V, the gel was transferred for western blotting using an iBlot (Invitrogen, US) and revealed after antibody incubations using a Fusion Solo (Vilber Lourmat, CH). Blot quantifications were done in ImageJ or Bio1D (Vilber Lourmat, CH).

### Crosslink experiments

Cells were washed two times with PBS, incubated 5 min at room temperature with crosslinker (1mM BMOE or 1mM DSS or 0.025% glutaraldehyde, Thermo Fisher, US), washed two times with PBS, lysed and boiled 5min at 95°C in non-reducing sample buffer before migration on 4–20% Tris-Glycine SDS-PAGE gels.

### Immunofluorescence

Cells were seeded on glass coverslips (N1.5, Marienfeld, D) for at least 48 h (including transfection). Fixation and permeabilization were performed either i) to preserve the secretory pathway as follows: cells were washed 3x with PBS and fixed with 3% paraformaldehyde for 20 min at 37°C, washed 3x with PBS, quenched with 50 mM of NH_4_Cl for 10 min at RT, washed 3x with PBS, permeabilized with 0.1% Triton X-100 for 5 min at RT and finally washed 3x with PBS; or ii) to preserve the cytoskeleton and ER membranes as follows: cells were washed 3x with PBS and fixed for 4 min at –20°C with precooled methanol and washed 3x with PBS. In both cases, the cells were then blocked overnight in PBS + 0.5% BSA (GE Healthcare, US). Immunostaining was performed as follows: the coverslips were incubated with primary antibody for 30 min at RT, washed 3x for 5 min with PBS – 0.5% BSA and incubated for 30 min at RT with secondary fluorescent antibodies (Alexa 488, 568 or 647, Invitrogen, US) and finally washed again 3x with PBS – 0.5% BSA prior to mounting in Mowiol. The coverslips were then imaged by confocal microscopy using a LSM710 microscope (Zeiss, D) with a 63x oil immersion objective (NA 1.4).

### Correlative Electron Microscopy

Cells were plated and transfected with ZDHHC6-GFP plasmids on glass coverslips coated in a 5-nm layer of carbon that outlined a numbered grid reference pattern. After 24 h, the cells were fixed for 60 min in a buffered solution of 2% paraformaldehyde and 2.5% glutaraldehyde at 25°C, and then washed 3x with cacodylate buffer. The coverslips were then mounted in a holder for fluorescence microscopy and the cells imaged using confocal microscopy (LSM700, Zeiss, 63x objective, NA 1.4). The cells of interest were imaged at a range of magnifications, and their location recorded according to the carbon grid pattern. The coverslips were then post-fixed with 1% osmium tetroxide and 1.5% potassium ferrocyanide in cacodylate buffer (0.1 M, pH 7.4) for 40 min at 25°C. After washing in distilled water and further staining with osmium alone followed by 1% uranyl acetate, they were dehydrated in a series of increasing concentrations of alcohol, then embedded in Durcupan resin, which was hardened overnight at 65°C. The next day, the resin containing the cells of interest was separated from the coverslips and mounted onto a blank resin block for ultrathin sectioning. Serial ultrathin sections were cut at 50 nm thickness and collected onto a formvar support film on single slot copper grids. Pictures were acquired at 80 kV using a transmission electron microscope (Tecnai Spirit, FEI Company, US).

### Focused Ion Beam Scanning Electron Microscopy (FIBSEM)

Cells of interest, recorded with fluorescent microscopy and prepared for electron microscopy (see above) were serially imaged using FIBSEM. Resin blocks were trimmed using an ultramicrotome so that the cell was located within 5 µm of the edge of the resin block. This block was then glued to aluminum stub, coated with a 20-nm layer of gold in a plasma coater, and placed inside the microscope (Zeiss NVision 40, Zeiss NTS). An ion beam of 1.3 nAmps was used to sequentially mill away 10-nm layers of resin from block surface to enable the cell to be serially imaged. Images were collected using the backscatter detector with the electron beam at 1.6 kV and grid tension set at 1.3 kV to collect only the highest energy electrons.

The final images were precisely aligned using the StackReg algorithm (56) in ImageJ, and the ER, mitochondria, nuclear membrane, and cell membrane segmented using the Microscopy Image Browser software (57). The mesh models were then exported to the Blender software (www.blender.org) for final rendering and visualization.

### Statistical analysis

Statistical analyses were performed using unpaired two-tailed Student’s t-test. Data are represented as means ± standard deviations. ns: not significant, \**p* < 0.05, \*\**p* < 0.01, \*\*\**p* < 0.001.

### Data availability

The authors declare that all data supporting the findings of this study are available within the paper and in the SI Appendix.

Further details on materials and methods can be found in SI Appendix.

## Acknowledgements

We thank Dr. Masaki Fukata for all of the ZDHHC-myc plasmids and ZDHHC16-FLAG; Dr. Ramanujan Hegde for the TRAPα antibody and Dr. Maurizio Molinari for the calnexin antibody. We thank the staff of the BioEM facility of EPFL, in particular, Marie Croisier and Stéphanie Clerc for their support regarding the EM experiments. The research leading to these results has received funding from the European Research Council under the European Union’s Seventh Framework Programme (FP/2007-2013)/ERC Grant Agreement n. 340260 – PalmERa’. This work was also supported by grants from the Swiss National Centre of Competence in Research (NCCR) Chemical Biology (to G.v.d.G) and the Swiss SystemsX.ch initiative evaluated by the Swiss National Science Foundation (LipidX) (to G.v.d.G and to V.H.).

